# Multi-omics biomarkers aid prostate cancer prognostication

**DOI:** 10.1101/2022.09.20.508244

**Authors:** Zhuoran Xu, Mohamed Omar, Elisa Benedetti, Jacob Rosenthal, Renato Umeton, Jan Krumsiek, Mark Pomerantz, Eddie Imada, Massimo Loda, Luigi Marchionni

## Abstract

Effective biomarkers and diagnostic tools are urgently needed in clinical settings for improved management of prostate cancer patients, especially to reduce over-treatment of indolent tumors and for early identification of aggressive disease. Gene expression signatures are currently the “gold standard” to provide guide clinical decision, however their clinical utility and interpretability is questionable. Multi-modal molecular profiling provides an holistic approach to systematically unravel the biological complexity underlying cancer pathogenesis, hence biomarkers developed using such an integrated approach hold the potential to more accurately capture cancer-driving alterations than signatures based on a single *omics* modality. Currently, however, robust and reproducible multi-*omics* biomarkers are still lacking for prostate cancer. In this study, we analyzed transcriptomics and metabolomics profiles jointly in a prostate cancer cohort and identified two prognostic signatures with high statistical powers (signature 1: EGLN3, succinate, trans-4-hydroxyprolin; and signature 2: IL6, SLC22A2, histamine). Our approach leveraged *a priori* biological knowledge of the cellular metabolism and gene circuitry, enabling the identification of dysregulated network modules. Functional bioinformatics analyses suggest that these signatures can capture relevant molecular alterations in prostate cancer tissues, including dysregulations of cellular signaling, cell cycle progression, and immune system modulation, stratifying patients in distinct risk groups. Next, we trained two gene expression signatures as a proxy for the multi-*omics* ones, extending our investigation to publicly available data, further confirming their prognostic values in independent patient cohorts. In summary, the analysis of multi-modal molecular grounded in cellular network biology represents a promising approach for the development of robust prognostic biomarkers of detecting and discriminating high grade disease.

## Introduction

Prostate cancer (PCa) is the most common malignant neoplasm of the urinary tract (Porzycki and Ciszkowicz, 2020; Litwin and Tan, 2017) and is a highly heterogeneous disease in almost every aspect from molecular diversity to clinical behaviour Barbieri and Shoag (2016); Carm et al. (2019). Most men diagnosed with prostate cancer have a favorable outcome of 99% overall ten-year survival if the disease is detected and treated early (Rebello et al., 2021). However, in some cases, the condition will metastasize having only a 28% five-year survival rate (Roviello et al., 2016). Thus, effective biomarkers are urgently needed to differentiate between indolent and aggressive prostate cancer to guide proper interventions.

Extensive efforts have been made to develop new prostate cancer diagnostic tools using different types of *omics* approaches (Doultsinos and Mills, 2021; Klein et al., 2014; Erho et al., 2013; Koo et al., 2019; Markert et al., 2011; Long et al., 2011). For example, Irshad et al. identified a three-gene expression signature (FGFR1, PMP22, and CDKN1A) that accurately predicts outcomes in low Gleason tumors (Irshad et al., 2013), while Yang et al. identified 3 metabolites (guanidinoacetate, phenylacetylglycine, and glycine) as potential novel biomarkers for prostate cancer detection and high grade disease discrimination (Yang et al., 2021). Another example is the identification of fifteen microRNAs recurrently associated with prostate cancer progression across multiple cohorts through a systematic review and reanalysis by Rana et al. (Rana et al., 2022) Although previous prostate cancer biomarker studies exhibited promising results, most relied on data from single *omics* technologies. Because the information offered by different data modalities (i.e., transcriptomics, genomics, epigenomics, metabolomics, etc.) does not necessarily overlaps, and often exhibits only modest correlation (Ghazalpour et al., 2011; Schwanhäusser et al., 2011), single modality biomarkers can only capture perturbations in specific domains, providing a skewed, and perhaps incomplete picture of the underlying biological processes.

On the other hand, as disease development involves complex interactions and alterations at multiple levels such as genome, epigenome, transcriptome, proteome, and metabolome, multi-*omics* profilings can describe biological processes comprehensively and systematically (Karczewski and Snyder, 2018). As shown in (Baranovskii et al., 2022), utilizing the transcriptome on top of gene panel features substantially improves drug response prediction performance in cancer. Therefore, biomarkers developed using multi-*omics* data can more accurately capture inter-patient heterogeneity than single-*omics* biomarkers (Olivier et al., 2019). As multi-*omics* profilings become more accessible, a few studies have started to utilize multi-*omics* profiles to develop predictive biomarkers for prostate cancer. For instance, Long et al. Long et al. (2011) identified a panel of 10 protein-coding genes along with two microRNA that are predictive of tumor recurrence from prostate cancer expression profiling. More recently, Wu et al. (Kiebish et al., 2020) combined proteomics, metabolomics, and lipidomics and identified a biomarker panel consisting of two proteins, one metabolite, and one phospholipid molecular that can predict biochemical recurrence in prostate cancer. However, integrating multiple types of *omics* profiles for biomarker development is not trivial and current studies mainly have two limitations. First, most studies analyzed profiles from different *omics* types in a sequential instead of simultaneous manner, limiting their ability to fully utilize the information from multi-*omics* data, especially the “cross-talks” between different *omics* types. Second, most previous studies provided a list of prognostic molecules instead of a decision rule for stratifying patients, undermining their potential translational values.

Since metabolites are the end products of upstream processes and carry information from both genetic and environmental changes (Guijas et al., 2018; Patti et al., 2012), integrating gene expressions with metabolite abundances will provide unprecedented opportunities to unravel biological complexities holistically. Thus, this study seeks to identify multi-*omics* signatures that can potentially be used as prognostic biomarkers by integrating patients’ transcriptomics and metabolomics profiles. We first constructed a multi-*omics* network that contains the minimal sets of aberrant gene-metabolite pairs, but it is sufficient to capture inter-patient heterogeneity using the Dana-Farber/Harvard Cancer Center (DF/HCC) cohort (Oh et al., 2006). We then identified two multi-*omics* signatures that can stratify prostate cancer patients into two risk groups with regard to disease-free survival. We further demonstrated that the identified signatures are associated with multi-faceted changes on pathways of signal transduction, cell cycle, and immune system. Finally, we constructed surrogate gene signatures based on patients’ recurrence risk groups, and extensively validated these in 8 external independent datasets as well as in the pooled dataset. In conclusion: (1) We identified two multi-*omics* signatures (signature 1: EGLN3, succinate, trans-4-hydroxyprolin; and signature 2: IL6, SLC22A2, histamine) that can accurately classify prostate cancer patients into two prognostic groups. (2) The surrogate genes signatures are also effective biomarkers, since the relative decision rule can be easily applied to other cohorts without the need of any re-calibration. (3) Our workflow paves the way for jointly analyzing multi-*omics* profiles for biomarker discovery.

## Results

### Study overview

We only included individuals with both transcriptomics and metabolomics profiles from the DF/HCC cohort (Oh et al., 2006) for biomarker development and functional analysis. The cohort includes 94 tumor samples, of which 48 have matched normal adjacency samples. Eighty-five patients have follow-up records with a median follow-up time of 2.02 years, including three lethal and eight progression cases. Summary statistics of demographic and clinical variables are provided in **Table 1**. In this study, we centered our analysis on gene-metabolite interactions. We first constructed a multi-*omics* covering network with a minimal set of aberrant gene-metabolite pairs to reduce noisy features from the data, but sufficient to account for observed inter-patient heterogeneity. Candidate signatures encompassing both genes and metabolites were then extracted based on the topology of the covering network, followed by survival analysis to asses their prognostic values (**Figure 1a**). After selecting prognostic multi-*omics* signatures, we conducted pathway analysis (using both gene expressions and metabolite abundances) to investigate pathological changes between the patients’ prognostic groups the signatures captured (**Figure 1b**). Finally, surrogate rank-based, single-*omics* gene signatures (based on the multi-*omics* classification results) were also obtained, to facilitate clinical usage when only gene expression data is accessible (**Figure 1c**).

**Table 1.** Descriptive statistics of DF/HCC cohort

|  |  | Overall<br>(N=94) |
| --- | --- | --- |
| **Age** | mean ± sd | 58.60 ± 7.17 |
|  | Missing | 3(3.2%) |
| **Race** | White | 77 (81.9%) |
|  | Black or African American | 9 (9.6%) |
|  | Other or Missing | 8 (8.5%) |
| **Hispanic** | Yes | 48 (51.1%) |
|  | No | 43 (45.7%) |
|  | Missing | 3 (3.2%) |
| **PSA before surgery** | median (25<sup>th</sup>, 75<sup>th</sup>) | 119(84.0, 174.0) |
|  | Missing | 13(13.83%) |
| **Gleason Risk** | Low (Primary Gleason ≤ 3) | 58 (61.7%) |
| | High (Primary Gleason $>$ 3) | 36 (38.3%) |

**Figure 1.**
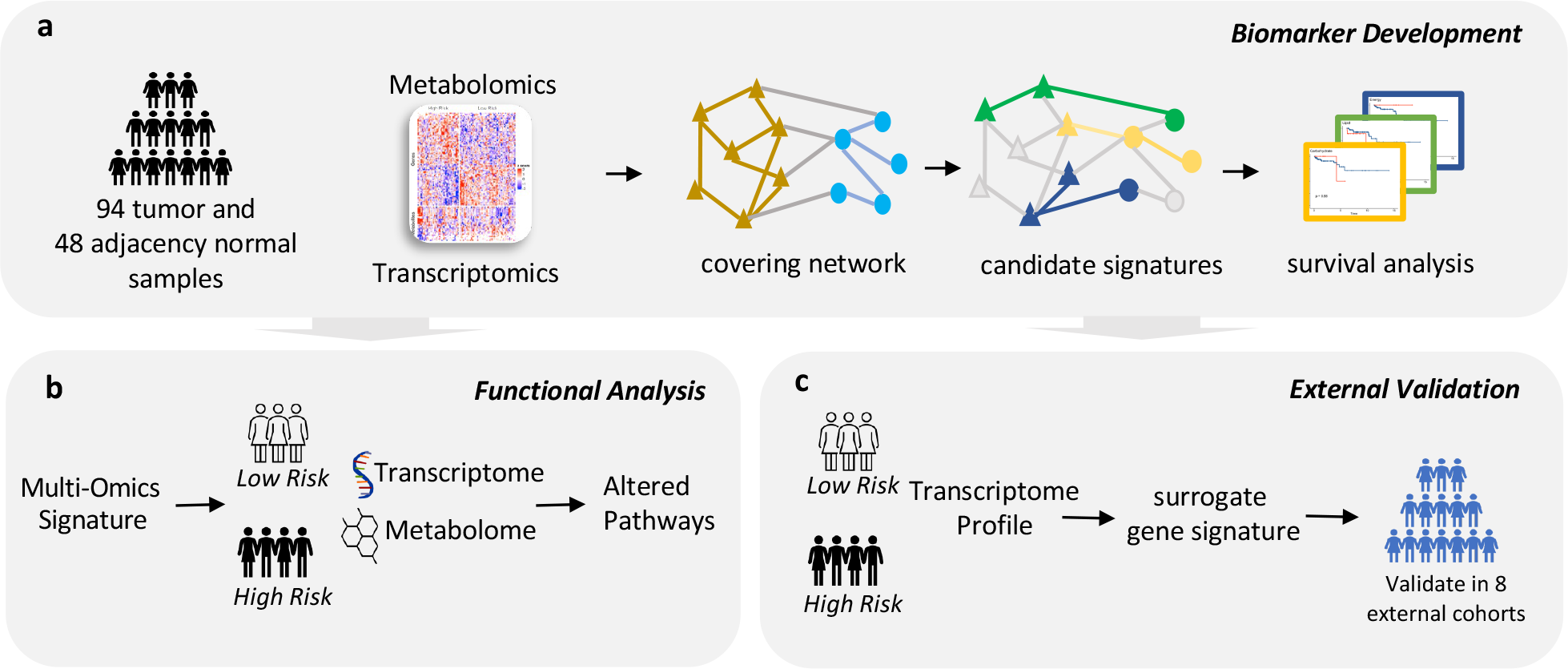
Study overview **a**. Biomarker development. 94 tumor and 48 adjacency normal samples with both transcriptomics and metabolomics profiles were used for developing prostate cancer biomarkers. A covering network that is parsimony and sufficient to account for inter-patient heterogeneity was constructed. Candidate multi-omics signatures were determined based on network topology followed by survival analysis to select prognostic signatures. **b**. Functional analysis. Pathway enrichment analyses were performed between patients’ risk groups using gene expression, metabolite abundances alone and jointly. **c**. External validation. Based on patients’ risk groups classified by multi-*omics* signatures, surrogate gene signatures using only gene expression data were trained and validated in external cohorts.

### Multi-*omics* covering network captures inter-patient heterogeneity efficiently

To capture systematic abnormalities that happen across different *omics* layers, we centered our analysis on paired gene-metabolite aberrations. We constructed the covering network followed as previously published (Ke et al., 2021) with some modifications. This approach was originally designed to select the minimal set of aberrant DNA-RNA pairs that yield a parsimonious sample-level representation rich enough to account for the observed inter-patient heterogeneity. Here we extended and applied the covering method to gene-metabolite profiles of prostate cancer. Briefly, genes (source level) and metabolites (target level) were first paired based on prior interaction knowledge from PathwayCommons (Rodchenkov et al., 2019) with the maximum length of the directed chain set as three. Thus, the initial network was comprised of 1,880,969 valid gene-metabolite pairs, involving 18,483 distinct genes and 111 distinct metabolites. For each gene-metabolite pair, a binary variable indicating whether the specific patient exhibits a “divergent” status compared to the normal range was derived using the divergence framework (Dinalankara et al., 2018, 2021). Next, we filtered out rare and independent gene-metabolite pairs based on binary divergence indicators, resulting in a network of 3,679 candidate pairs accounting for 12,245 unique genes and 71 unique metabolites. After the optimization procedure, 20 gene-metabolite pairs covering 98.9% of the patients in the cohort with maximum divergence probability were selected to obtain the final covering network. We filled all intermediary genes between each selected source-target pair and all within- and between-*omics* interactions based on PathwayCommons. The final, complete covering network consisted of 20 source genes, 34 intermediary genes, 12 target metabolites, 117 gene-gene interactions, 1 metabolite-metabolite interaction, and 30 gene-metabolite interactions (**Figure 2a**).

**Figure 2.**
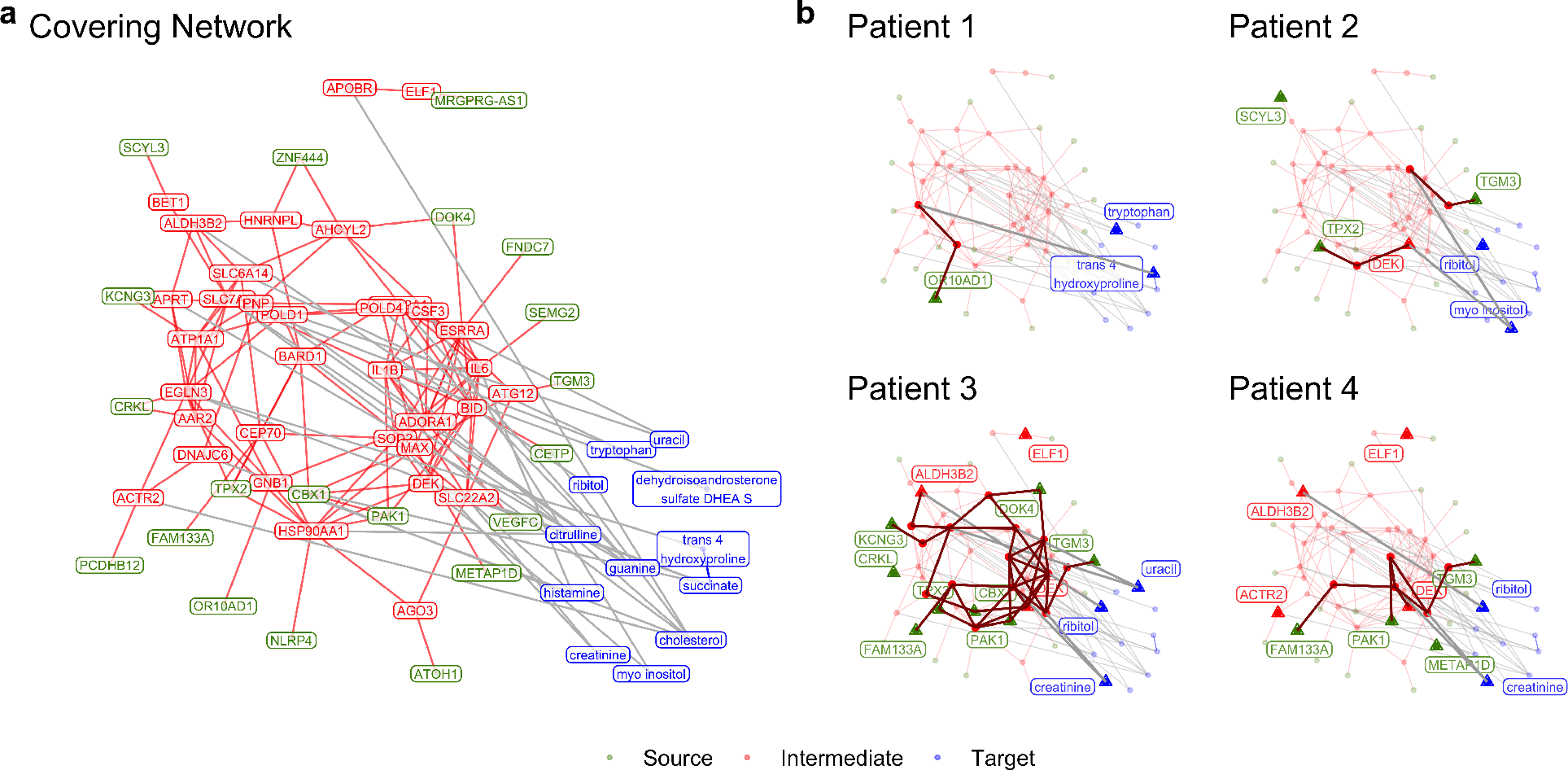
Covering network. **a**. Complete network covering the whole cohort. **b**. Different patients exhibited distinct aberrant patterns captured by network.

Since prostate cancer is a heterogeneous disease, an efficient cohort-level representation should be not only parsimonious but also sufficient to cover the variance of cohort patients from different angles. Accordingly, different patients should exhibit distinct molecular aberrations on the covering network, and at the same time, different source-target pairs can stratify different subsets of patients. In other words, a covering network is the desired representation if similarities of both inter-pair (parsimony) and inter-patient (sufficient) are relatively low. We then examined patients’ divergence status of every gene, metabolite, and gene-metabolite pair in the covering network to inspect the inter-patient heterogeneity captured. It can be seen from the results that different patients exhibited distinct molecular aberration patterns in the covering network (**Figure 2b**, **Figure S1**). We also quantitatively evaluated inter-patient similarities based on the divergence status of 20 gene-metabolite pairs and inter-pair similarities based on 94 patients’ divergence status. Results showed that both inter-patient and inter-pair similarities are numerically low, with average Jaccard coefficients of 0.050 and 0.024, respectively. All above results demonstrated the multi-*omics* covering network we constructed can efficiently capture observed inter-patient heterogeneity.

### Two multi-*omics* signatures are prognostic

We aimed to identify prognostic multi-*omics* signatures from the covering network. Candidate signatures were first determined by a diffusion process based on network topology by assuming that topologically closed molecules are also functionally related. Each connected gene-metabolite pair was set as seeds to enforce candidate signatures containing cross-modal information. Nineteen candidate signatures were obtained from the complete covering network, with the total number of genes and metabolites varying from 3 to 8 (**Figure S2**). We then stratified the patients into low and high risk groups for each candidate signature by performing a hierarchical clustering using the corresponding molecules’ abundances. By comparing disease-free survival between patient risk groups, two prognostic multi-*omics* signatures (signature 1: EGLN3, succinate, trans-4-hydroxyproline, Log-Rank test: p=0.019; and signature 2: IL6, SLC22A2, histamine, Log-Rank test: p=0.017) were identified from the covering network (**Figure 3**). Because the patient number of the cohort is limited, we grouped patients with primary Gleason scores of 3 into the low Gleason risk group and the rest into the high Gleason risk group (**Table 1**). After stratifying patients into high- and low-Gleason groups, we found signature 1 is significantly prognostic in high-Gleason risk group (Log-Rank test: p=0.0073) and signature 2 is significantly prognostic in low-Gleason group (Log-Rank test: p=0.0073). We then created an ensemble signature that categorized patients into the high-risk group if they were high-risk patients by both signatures. Such signature ensemble can successfully identify patients who relapsed in the whole cohort patients (Log-Rank test: p<0.0001) as well as in a stratified analysis by Gleason score (Log-Rank test: high-Gleason group: p=0.0011; low-Gleason group: p<0.0001). To test whether the signatures can add additional prognostic values on top of widely used Gleason scores and Prostatic Specific Antigen (PSA), we conducted a likelihood ratio test by nesting multivariate Cox models with and without the signature risk predictions, after adjusting for Gleason risk groups and PSA. Results demonstrated that adding signature 1 or the signature ensemble on top of Gleason and PSA can add accuracy in predicting patients’ of disease-free survival (Likelihood ratio test: signature 1: p=0.019; signature ensemble: p=0.0057), while signature 2 didn’t exhibit any significant improvement(Likelihood ratio test: p=0.615). Finally, we also conducted permutation analysis to circumvent the possibility that the signatures were determined by randomness, ensuring statistical power. The identified signatures have statistical powers of 95.2% (signature 1) and 94.5% (signature 2), respectively.

**Figure 3.**
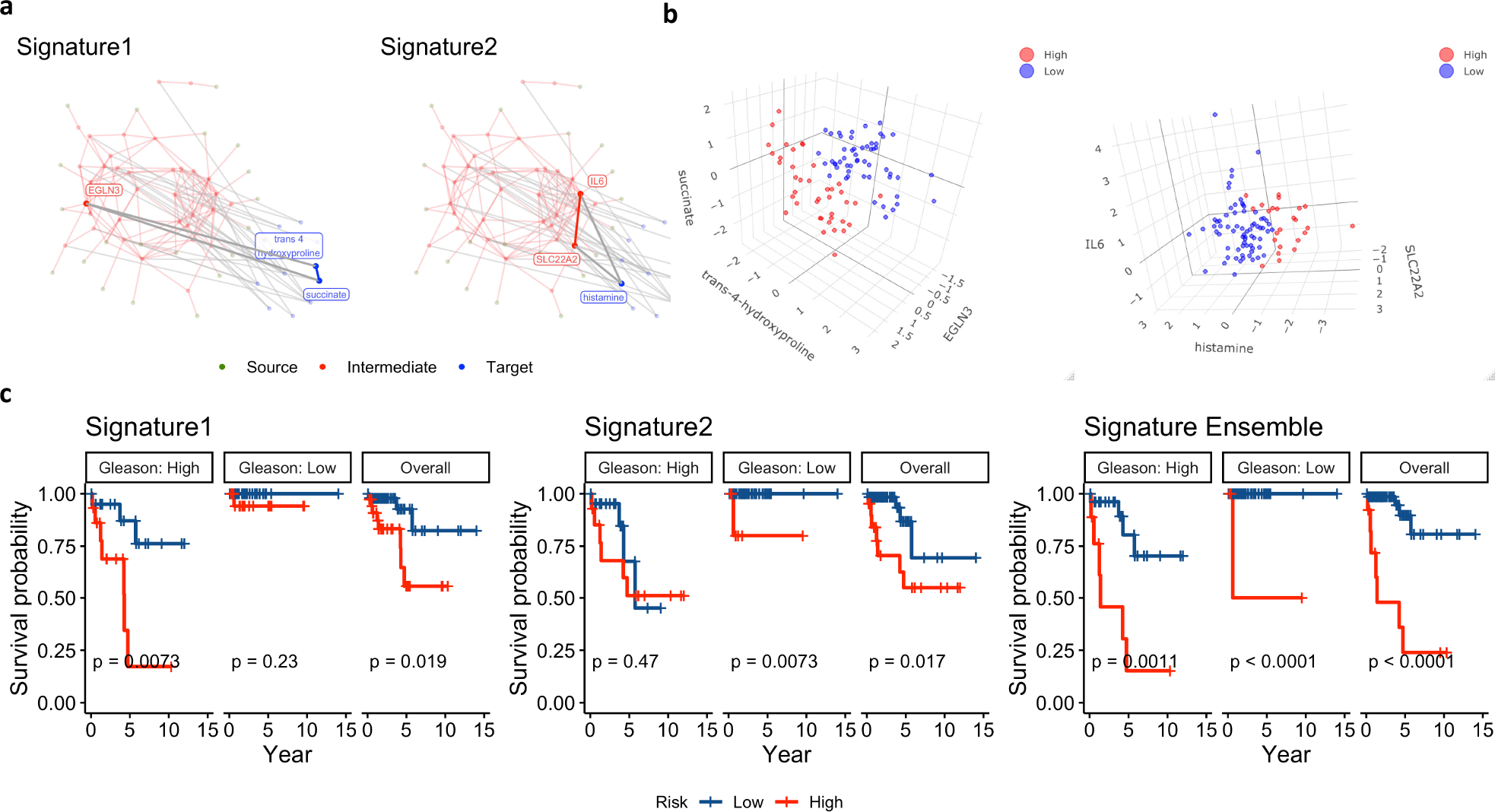
Two prognostic multi-*omics* signatures were identified from the covering network. **a**. Both of two multi-*omics* signatures identified from the covering network contain gene(s) and metabolite(s). **b**. Two signatures stratified patients into two risk groups viewed in 3D space. **c**. Kaplan-Meier curves between patient risk groups with and without stratification by Gleason scores. Signature 1 is prognostic in high Gleason risk group and signature 2 is prognostic in low Gleason risk group. Signature ensemble is prognotic in both Gleason risk groups.

### Patients’ prognostic groups associate with profound molecular alterations

To investigate underlying pathway alterations between high- and low-risk groups captured by the two multi-*omics* signatures, we performed enrichment analyses on single- and multi-*omics* levels. Specifically, we performed differential genes and metabolite analysis between the prognostic groups and used the results as input for pathway enrichment exploration. For the joint genes/metabolites enrichment, we performed a combined pathway analysis capturing biological changes across *omics* layers (Canzler and Hackermüller, 2020). For signature 1, we found 1,151 genes and 28 metabolites with significant differences between low and high-risk prognostic groups. Likewise, we found 728 differential genes and 12 differential metabolites between prognostic groups defined by signature 2. Multi-faceted pathway changes were identified for both signatures, including gene sets involved in signal transduction, metabolism of proteins, and immune system modulation. We noticed that signature 1 captured perturbations of activated pathways relating to signal transduction on both gene and metabolite levels. However, signature 2 primarily identified pathway changes on the gene level, such as significantly activated DNA replications and cell-cycle-related pathways. In addition, patients classified into the high-risk group by signature 1 had activated interleukin signaling pathways, while high-risk patients identified by signature 2 showed a broader scope of up-regulated related to immune functions, including cytokine signaling, adaptive immunity, and the innate immune system.

We then conducted cell type enrichment deconvolution analysis using bulk gene expression data to compare cell type differences between patient prognostic groups. We found that high-risk patients in signature 1 had more regulatory T cells (p=0.016) when compared to low-risk patients. While high-risk patients in signature 2 had elevated levels of B cells (p=0.039) and macrophages (p=0.039), specifically M2 macrophages (p=0.030) compared to low-risk patients. Finally, we compared sample-level signature scores between prognostic groups to evaluate the associations between existing prognostic prostate cancer signatures with our multi-*omics* signatures. Results suggested that signature 1 high-risk patients had significantly higher signature scores for a 28-gene hypoxia-related signature developed by Yang et al. (Yang et al., 2018) (p=0.014), confirming the role of EGLN3 in hypoxia (Strocchi et al., 2022). However, high-risk patients of signature 2 exhibited low signature scores for Talantov(Talantov et al., 2010) (p = 0.005) and Planche (Planche et al., 2011) signatures (p=0.035).

### Surrogate gene signatures are prognostic in external validation cohorts

Since multi-*omics* data sets are relatively rare, we we constructed single modality gene signatures aiming to proxy our multi-*omics* signatures, with the ultimate goal of facilitating clinical applications when only gene expression data is available. We used pairwise gene comparisons (Geman et al., 2004; Marchionni et al., 2013; Afsari et al., 2014) as binary features and fit lasso regression models to train surrogate gene signatures, which is robust to quantization effects and invariant to pre-processing. Patients’ prognostic classifications by our multi-*omics* signatures were used as labels. The whole DF/HCC cohort was used as training dataset, where surrogate gene signature 1 and signature 2 achieved accuracy of 86.17% and 94.68% (**Figure 4a**), respectively. To ensure these surrogate gene signatures still held prognostic value, we performed log-rank tests and found surrogate gene signatures are still prognostic (**Figure 4b**; surrogate signature 1: p=0.037, surrogate signature 2: p=0.024, surrogate signature ensemble: p <0.0001). Consistent with corresponding multi-*omics* signatures, surrogate gene signature 1 is prognostic for high Gleason patients (log-rank p=0.023) and surrogate gene signature 2 is prognostic for low Gleason patients (log-rank p=0.016). However, no evidences can show any surrogate gene signatures can statistically add propensity on top of Gleason risks and PSA values (Likelihood ratio test: surrogate signature 1: p=0.062; surrogate signature 2: p=0.836; surrogate signature ensemble: p=0.061). We then validated the surrogate gene signatures in 8 independent external cohorts as well as in the resulting pooled cohort. Surrogate signature 1 and signature 2 are prognostic in 2 and 4 cohorts, respectively. The surrogate ensemble signature is more robust and prognostic in 5 out of 8 cohorts. All surrogate signatures are prognostic in the pooled cohort (**Figure 5a**). We finally performed log-rank test adjusting for Gleason score. Surrogate gene signatures are prognostic in 1 and 2 out of 8 cohorts, respectively, while the signature ensemble is prognostic in 3 cohorts, but not the pooled cohort (**Figure 5b**).

**Figure 4.**
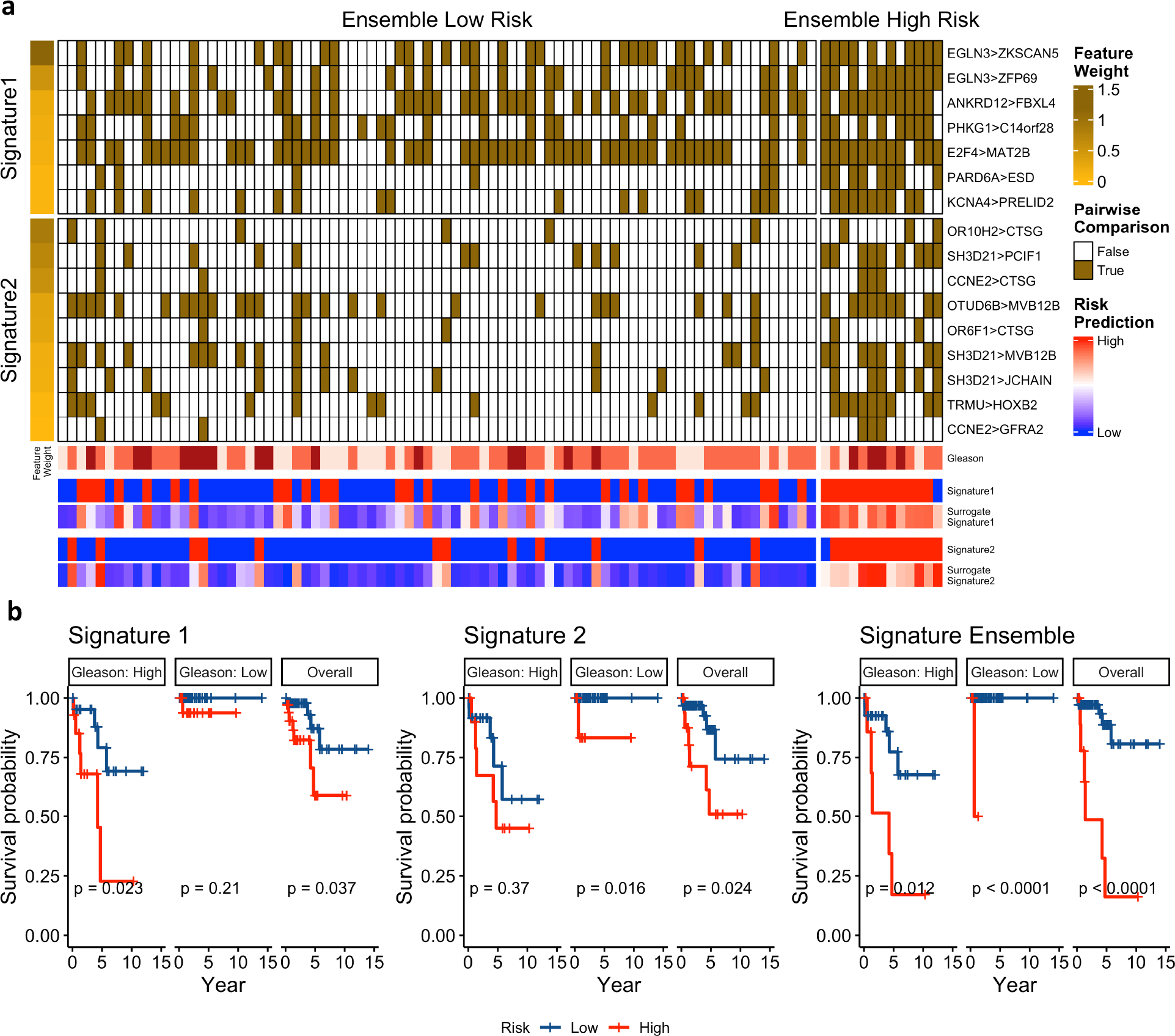
Surrogate gene signatures trained in DF/HCC cohort as multi-*omics* signature proxies. **a**. Heatmap of surrogate gene signatures with each column of the heatmap body representing one patient and each row representing one pairwise gene comparison. Heatmap body was split by predictions of surrogate signature ensemble. Annotations of the heatmap indicate original labels of patients’ risk group of two multi-*omics* signatures and predicted risk scores by two surrogate gene signatures **b**. Kaplan-Meier curves between patient risk groups classified by surrogate gene signatures and surrogate gene signature ensemble with and without stratification by Gleason scores.

**Figure 5.**
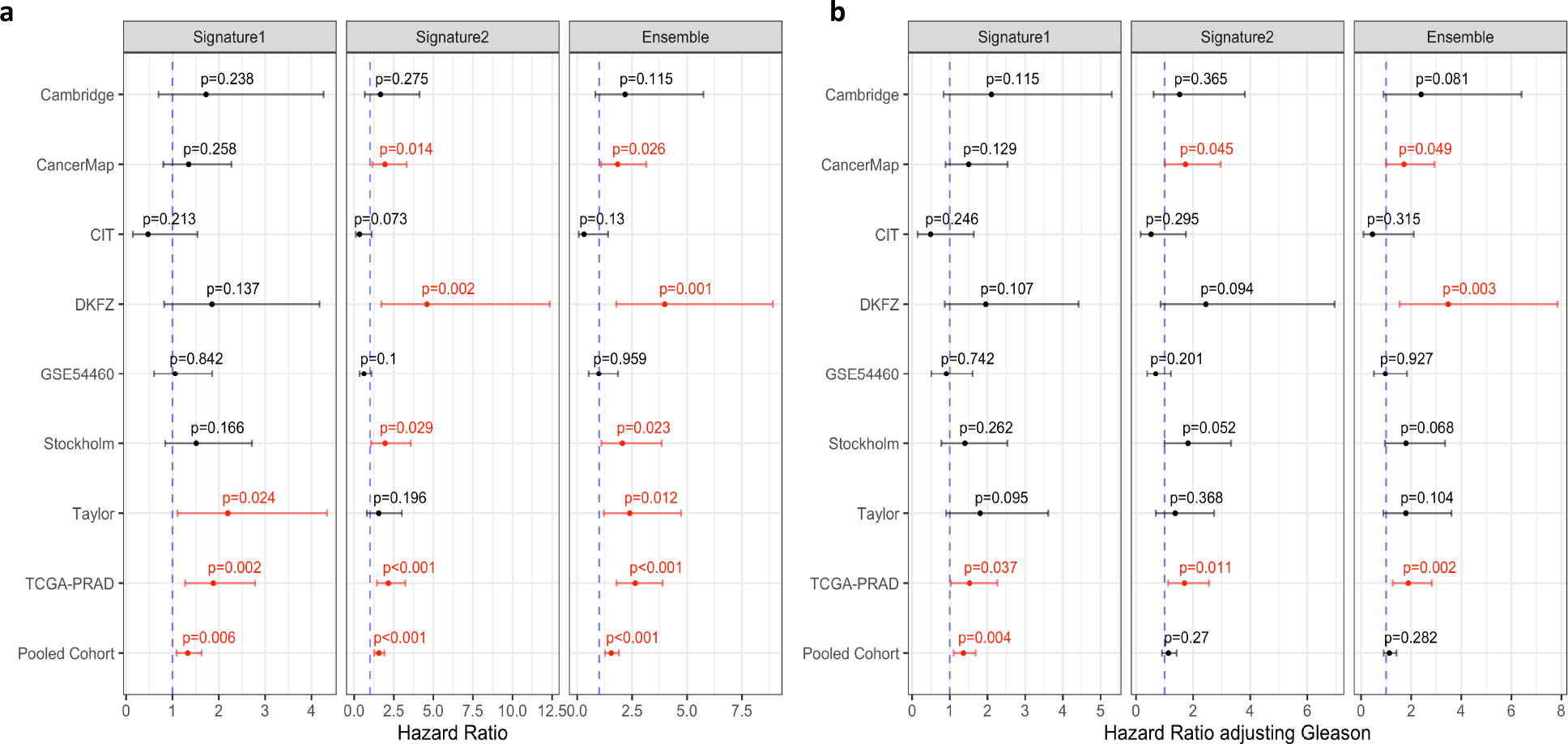
Validation of surrogate gene signatures in external prostate cancer cohorts. **a**. Surrogate gene signatures as the single predictor. **b**. Surrogate gene signatures adjusting for Gleason score. Plots show hazard ratios of high risk groups classified by each surrogate signature. Red indicate significance. Error bars represent 95% confidence interval of hazard ratios.

## Discussion

Extensive efforts have been made to develop prostate cancer molecular prognostic biomarkers using *omics* profiles to differentiate between indolent and aggressive disease. However, most existing signatures rely on a single type of feature, such as gene expression, and only capture a partial view of inter-patient heterogeneity. Because prostate cancer is highly heterogeneous, using single-*omics* profile to develop predictive biomarkers might be insufficient to recapitulate systematic alterations related to the disease for accurate predictions (Olivier et al., 2019). Therefore, integrating multiple profiles, such as gene expressions and metabolite abundances, and especially their cross-talks, offers exceptional opportunities for advancing diagnosis, monitoring, and treatment of prostate cancer.

Hance, we analyzed gene-metabolite interactions and used multi-*omics* profiles to identify prognostic signatures for prostate cancer. We first constructed a covering network to account for inter-patient variability (**Figure 2, S1**). This covering network can be interpreted as a feature selection step, where only important, but not redundant features, reflecting gene-metabolite interplay, are kept in the network. Thus, the sub-networks within the covering network are expected to reflect inter-patient molecular heterogeneity. We then dissected such covering network into candidate signatures, identifying those with prognostic values. We utilized the topological structure of the covering network to select candidate signatures, assuming topologically closed molecules are more likely to be functionally related (**Figure S2**). With this approach, we identified two prognostic signatures, containing both genes and metabolites, that can accurately stratify patients into high- and low- recurrence risk groups (**Figure 3a-b**, signature 1: EGLN3, succinate, trans-4-hydroxyprolin; and signature 2: IL6, SLC22A2, histamine). The EGLN3 gene in the first signature have long been known to relate with hypoxia, a cancer hallmark(Schödel and Ratcliffe, 2019), while succinate is a key metabolic factor in the cancer-immune cycle (Jiang and Yan, 2017), and an initiator in tumorigenesis and cancer progression (Zhao et al., 2017). In the second signature, IL6 has also been shown to be involved in prostate cancer initiation, invasion and metastasis (Culig and Puhr, 2018). We further performed permutation studies to ensure the robustness of identified signatures, and demonstrated high statistical powers (95.2% for signature 1 and 94.5% for signature 2, respectively). Although the two multi-*omics* signatures are both prognostic in the DF/HCC cohort, we noticed their different roles in patients of different Gleason groups. Specifically, Signature 1 is prognostic mainly in patients with high-Gleason score, while Signature 2 is prognostic primarily for patients with low-Gleason score (**Figure 3c**). And the signature ensemble was shown to be prognostic in both Gleason patient groups.

To elucidate functional differences between high- and low-risk patient groups captured by these signatures, we conducted gene, metabolite, and multi-*omics* pathway enrichment analysis (**Additional file**). This approach revealed several key biological processes that are differently represented in distinct prognostic groups. For instance, the high-risk group defined by signature 1 has many up-regulated pathways involved in immuno-modulation, including IL10 signaling, which inhibits the activation of antigen-presenting cells (APCs) like macrophages and dendritic cells (DCs) and induces TH2 cells differentiation from naive CD4+ T cells(Ouyang and O’Garra, 2019). Similarly, the high-risk group also showed up-regulated IL4 and IL13 signaling pathways, which are both known to contribute to cancer progression and are associated with poor prognosis (Barderas et al., 2012; Roca et al., 2012). On the other hand, the high-risk group was also found to have down-regulation of molecular processes involved in DNA damage repair, including the Global Genome Nucleotide Excision Repair (GG-NER) and the Transcription-Coupled Nucleotide Excision Repair (TC-NER) pathways (Lockett et al., 2005). Finally, the high-risk group identified by signature 2 showed up-regulation of different molecular and biological processes. Specifically, we were able to reveal the up-regulation of the pathways involved in the regulation of RUNX2 and RUNX3 expression and activity. RUNX2 has been shown to play an important role in mediating prostate cancer bone metastasis, especially in tumors with PTEN loss (Zhang et al., 2011). RUNX3 has a tumor suppressor role and its loss is associated with tumor progression (Ashe et al., 2021). Interestingly, the ABC transporter disorders pathway was also found up-regulated in the high-risk group predicted by signature 2, a biological process linked to chemoresistance and potentially worse survival in this group (Robey et al., 2018). Taking together, multi-*omics* signatures can capture patients’ multi-faceted alterations associated with tumor progressions in a systematical way.

Unfortunately, the availability of prostate cancer datasets with paired transcriptomic and metabolomic data is scarce. To overcome this issue, we constructed single-*omics* signatures as proxies to predict patients’ risk groups resulting from the multi-*omics* signatures. Hence, we trained two surrogate gene expression signatures in the DF/HCC cohort, and then validated these externally. Most decision rules built on high-throughput *omics* data use continuous expression or abundance values can suffer from a lack of robustness and low cross-study reproducibility, limiting their translational potential. Here we adopted a rank-based approach for constructing surrogate signatures so that they are robust to quantization effects and invariant to pre-processing. Although the surrogate gene signatures achieved high performance in the DF/HCC cohort (**Figure 4**), they were intrinsically different from the multi-*omics* signatures due to the lack of integrated, cross-*omics* information. Nevertheless, we found that these surrogate signatures exhibited prognostic values in 5 and 4 out of 8 external datasets before and after adjusting for Gleason risk groups (**Figure 5**).

Overall, we identified two multi-*omics* signatures through the joint analysis of cross-*omics* interactions from transcriptomic and metabolomic profiles. The two signatures we identified can capture multi-faceted pathway alterations within prostate cancer patients in the DF/HCC cohort, and they were shown to be prognostic. These surrogate gene signatures, especially after ensembling, are still effective biomarkers when validated in external, independent cohorts. While further functional validation of these multi-*omics* signatures are still needed, we believe this work highlights the importance of integrating multi-*omics* profiles for biomarker discovery and development.

## Methods

### Patient Cohorts

All samples used for biomarker development and functional analysis are from the Dana-Farber/Harvard Cancer Center Cohort (DF/HCC) as described previously (Oh et al., 2006). The DF/HCC Cohort includes patients’ clinical information, blood samples, and tissue biopsy samples from more than 4,000 prostate cancer patients. Each subject consented to the use of clinical data and specimens for research purposes. In this study, we only included subjects with both transcriptomics and metabolomics profiles from fresh-frozen radical prostatectomy specimens, where 48 patients having both tumor and adjacency normal samples and 46 patients only having tumor samples. Among all 94 patients, 85 patients have follow-up records with a median follow-up length of 2.02 years, including three lethal and eight progression cases. Since there is no publicly available prostate cancer cohorts with both transcriptomics and metabolomics profiles, we built surrogate single-*omics* signatures for validating proposed biomarkers externally using transcriptomics datasets from PCaDB (Li et al., 2021). All tumor samples from the cohorts that have follow-up information were included for testing surrogate gene signatures except for CPC-Gene and Belfast cohort due to half of the gene pairs not being found in the data. In total, we used 8 external validation cohorts, including CancerMap, GSE54460, Taylor, Cambridge, CIT, DKFZ, Stockholm and TCGA-PRAD. We also pooled all above cohorts as a heterogenuous pooled cohort (**Additional file1**).

### Construction of a covering Network

Before constructing the covering network, we derived divergence statistics (Dinalankara et al., 2018), a binary random variable, for every tumor sample using the `Divergence` (Dinalankara et al., 2021) R/Bioconductor package to indicate whether each gene expression and metabolite abundance is aberrant compared to the baseline range calculated based on 48 normal samples. We define a gene-metabolite pair as divergent if the divergence statistics of the gene is correlated with that of the metabolite using Chi-squared test. Prior-knowledge interaction information between gene-gene, gene-metabolite, and metabolite-metabolite was obtained from PathwayCommons(Rodchenkov et al., 2019). We only kept 18,490 genes and 111 metabolites that can be found in both *omics* profiles and PathwayCommons for the subsequent analyses. Except for known gene-metabolite pairs, we also included the pairs if the source gene can get to the target metabolite within three steps with only passing through other genes, aiming to capture potential gene-metabolite indirect interactions based on current network knowledge. For example, we can match three gene-metabolite pairs from the chain of *gene*_1_ − *gene*_2_ − *gene*_3_ − *metabolite*. 1,880,969 valid gene-metabolite pairs with 18,483 distinct genes and 111 distinct metabolites were then obtained. The Chi-square test was used to test if the divergence status of source genes and target metabolites within pairs are dependent. All 1,847,998 independent pairs with p-value larger than 0.05 were filtered out. 29,292 rare pairs with divergence probability less than 2% were also excluded. After the above filtering procedures, 3,679 candidate pairs with 12,245 unique genes and 71 unique metabolites were left for optimizing the covering network. Finally, twenty gene-metabolite pairs covering 98.9% of the cohort patients with maximum divergence probability were selected. And the entire covering network consisted of 20 source genes, 34 intermediary genes, and 12 target metabolites. We also connected the molecules if they have a known interaction based on PathwayCommons and finally got 117 gene-gene interactions, 1 metabolite-metabolite interaction, and 30 gene-metabolite interactions for the covering network. Inter-patient Jaccard similarity was evaluated by

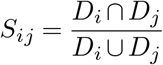

where *S_ij_* denotes similarity between patient *i* with patient *j*, and *D_i_* refers the set of divergence pairs of patient *i*. Similarly, inter-pair Jaccard similarity was evaluated by

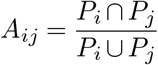

where *A_ij_* denotes similarity between the *i^th^* gene-metabolite pair with the *j^th^* gene-metabolite pair, and *P_i_* refers the patient set with *i^th^* gene-metabolite pair being diverged.

### Identification of prognostic multi-*omics* signatures

A diffusion procedure was performed to identify candidate signatures of which molecules are topologically close in the covering network. Because the regulatory directions are not important here, we consider the covering network as an undirected graph *G* = (*V, E*), where *V* is the set of nodes and *E* is the set of edges. We denote the adjacency matrix of *G* as *W*, where *W_i,j_* = 1 if there is an edge between node *i* and node *j*, otherwise as 0. 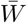 is column-normalized matrix of *W* to ensure convergence. Since our analysis focused on integrating both *omics* types, we forced selected signatures to contain both gene(s) and metabolite(s) by setting starting seeds from each connected gene-metabolite pair. We denote *P*^(0)^ as seed matrix with shape of 66 × 30, where the entry of 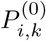 representing whether the *i^th^* node in the covering network belonging to the *k^th^* connected gene-metabolite pair. 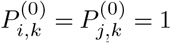 if node *i* and node *j* are connected with one being gene another being metabolite, and other entries of the *k^th^* column in *P* are 0. We then iteratively diffuse *P*^(0)^ on 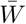 with *α* (0 ≤ *α* ≤ 1) controlling for the balance between local and global network similarities. Such that,

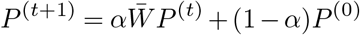

When *α* approaches 0, more local similarities will be used; otherwise, more global network information will be involved. We set *α* = 0.75, based on previous studies (Ruan and Wang, 2020). *t* represents iteration step, and we consider convergence if || *P*^(*t*+1)^ − *P*^(*t*)^ ||_1_ < 10^*−6*^. After the diffusion processes, we can find some gene-metabolite pairs, especially the ones connecting to the same molecules, have similar diffused score vectors (column vectors of *P*^(*t*)^, see **Figure S2a**) that can be used as gene-metabolite pair representations. We then performed hierarchical clustering of the 30 edges connecting gene-metabolite pairs using Ward’s minimum variance method and identified edge clusters (**Figure S2b-c**). Molecules connected by the edges from the same cluster will be treated as one candidate signature. Thresholds of 0.45, 0.65, and 0.85 were set to prune the hierarchical tree, from where most of the edges can cluster with at least one other edge to no single edge was left out **Figure S2c**.

We employed patients’ *omics* profiles that were scaled into continuous z-scores for risk stratification, which was achieved by hierarchical clustering using Ward’s minimum variance and complete linkage method. We then performed a log-rank test to compare disease-free survival times between two groups classified by each signature and only left prognostic signatures with a p-value<0.05. We also excluded the signatures if the smaller patient cluster had less than 10% of the patients. We then permuted the abundance of all features of each prognostic signature and repeated hierarchical clustering and survival analysis 1,000 times. The statistical power for the selected signature was calculated by the percentage of getting p-values > 0.05 in permutation studies.

### Construction of surrogate signatures

Surrogate gene signatures were all trained in DF/HCC cohort and validated in independent external test datasets as mentioned above. For each multi-*omics* signature, patients’ risk labels were used to train the surrogate gene signature classifier. The procedures are as following: (1) Differentially expressed genes between two patient risk groups (t test: p values<0.01) were selected as context genes. (2) Within context genes, we only kept the genes that are significantly correlated with at least one of the metabolites in the multi-*omics* signature with p values less than 0.01. (3) If genes in the multi-*omics* signatures were not in the gene list, we added those genes into the genes. (4) The left genes were paired with one being up-regulated and the other being down-regulated. (5) We then derive pairwise score (Geman et al., 2004) for each gene pair given by:

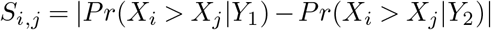

*S_i,j_* denotes gene pair score for gene *i* and gene *j*, where gene *i* is the up-regulated gene and gene *j* is the down-regulated gene.*X_i_* and *X_j_* are gene expression values for gene *i* and *j*. *Y*_1_ and *Y*_2_ are risk group labels. (6) To further reduce the complexity and noises, we only kept top 50 gene pairs with highest scores to training the surrogate classifier. Thus each patient has 50 binary indicator variables representing whether gene *i* has a higher expression level than gene *j*. (7) 5-fold cross-validation lasso logistic regression was then performed to select important gene pairs and derive parameter weights for each pair. Predictions of lasso regression models were used to classify patients into different risk groups. 0.5 was used as the cutoff for the training set. Each external validation cohort used the corresponding median value as the cutoff.

### Statistical analysis and data availability

All statistical and bioinformatics analyses were performed in R, version 4.0.3. Chi-square or Fisher’s exact test was performed as appropriate for selecting significant associated gene-metabolite pairs for constructing the covering network and examining associations between patient risk groups with genotype and Gleason scores. Log rank test was used for comparing Kaplan-Meier survival curved between high- and low-risk patient groups stratified by different signatures, where time to any events including biochemical recurrence, metastasis and death was used as composite survival endpoints. Gene set enrichment analysis and metabolite set enrichment analysis were conducted using `fgsea` (Korotkevich et al., 2016) R package and Reactome pathway database (Gillespie et al., 2021). Gene-metabolite multi-*omics* pathway enrichment analysis were performed followed the workflow proposed by Canzler et al (Canzler and Hackermüller, 2020). through combining p-values from each of single-*omics* enrichment analysis using Stouffer’s method. Unless explicitly mentioned, otherwise all statistical tests were two-sided, with p<0.05 indicating significance. Data used in the manuscript can be found at https://github.com/Karenxzr/MultiModalPC.

## Supplementary Information

**Figure S1.**
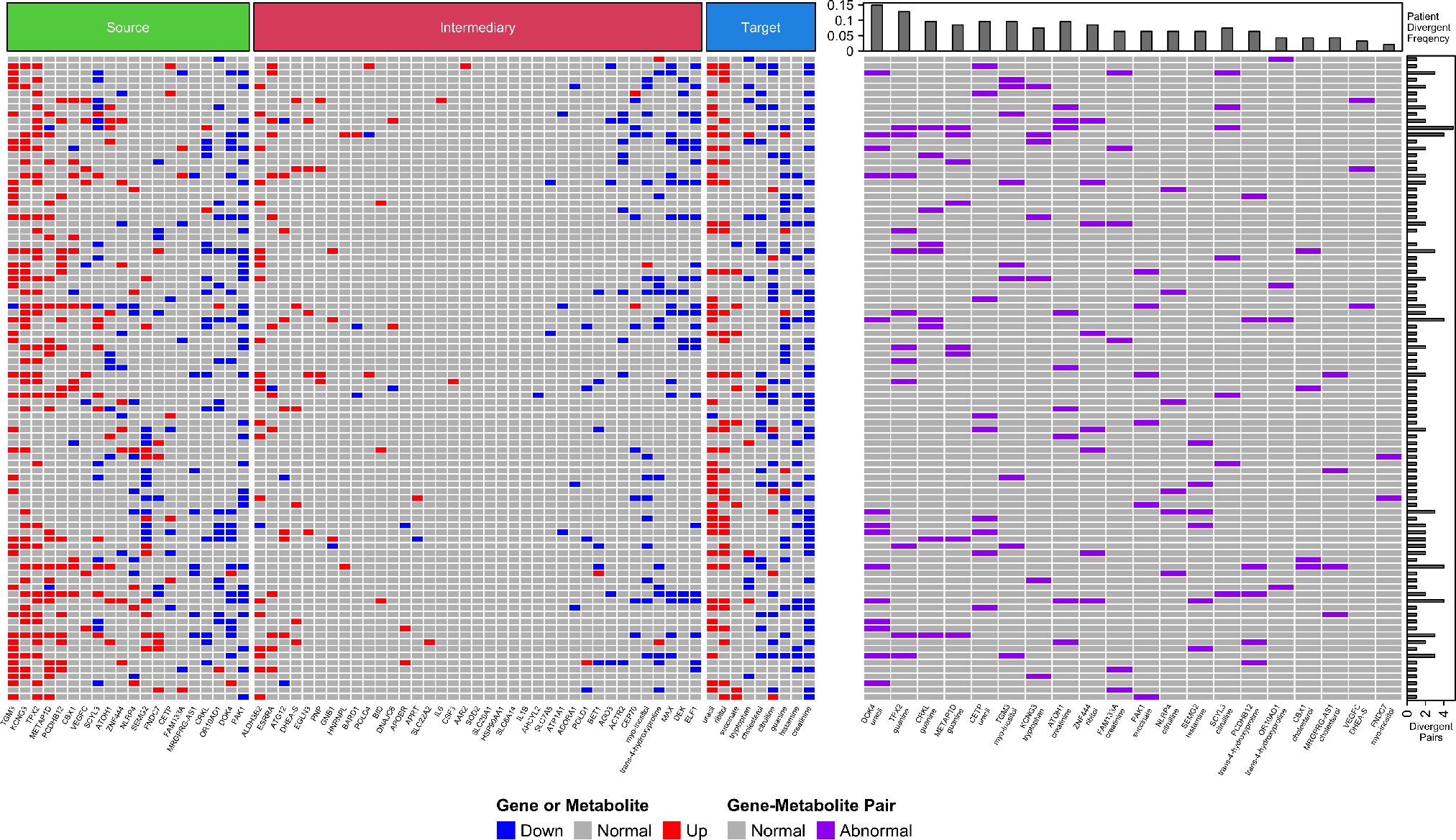
Inter-patient heterogeneity captured by the covering network. (**a**) Patients’ divergent status on single molecular level of each gene and metabolite in the covering network. (**b**) Patients’ divergent status of gene-metabolite pairs in the covering network.

**Figure S2.**
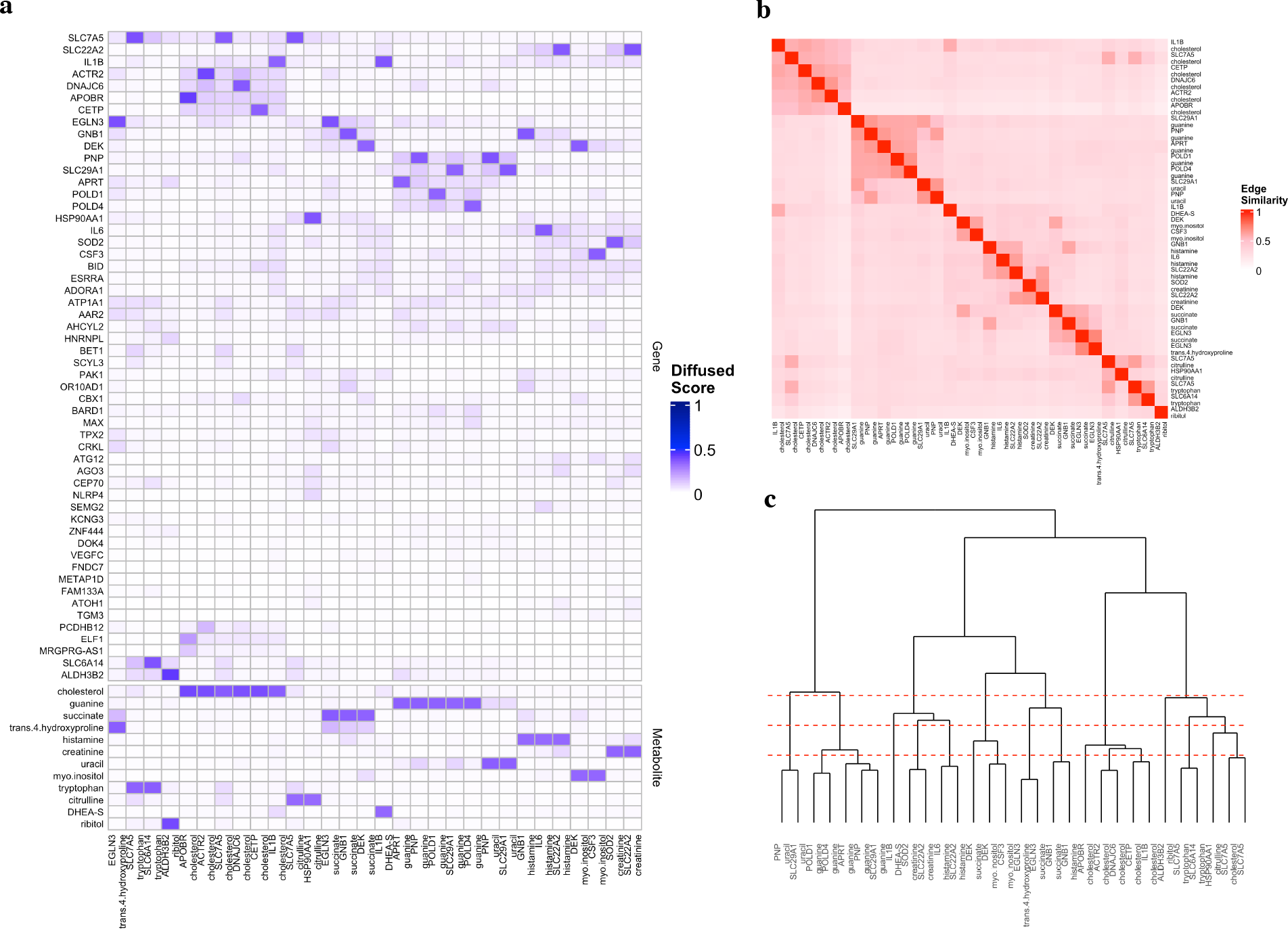
Identification of candidate signatures from the covering network. (**a**) Diffused scores on the covering network from every gene-metabolite pair. Each column represents seed gene-metabolite pairs. Each row represents scores of each gene and metabolite on the covering network after diffusion. (**b**) Similarity matrix of gene-metabolite pairs of the covering network. (**c**) Hierarchical clustering with different thresholds to decide candidate signatures composed of similar gene-metabolite pairs.

### Additional file1

Sheet1-2: Gene set enrichment analysis results for two multi-omics signatures. Sheet3-4: Metabolite set enrichment analysis results for two multi-omics signatures. Sheet5-6: Multi-omics enrichment analysis results for two multi-omics signatures. Sheet7: Summary statistics of external validation cohorts. Sheet8: Patient counts by signature risk groups and Gleason risk groups

