## Supplementary figures and images for "Multi-omics biomarkers aid prostate cancer prognostication"

### Figure S1

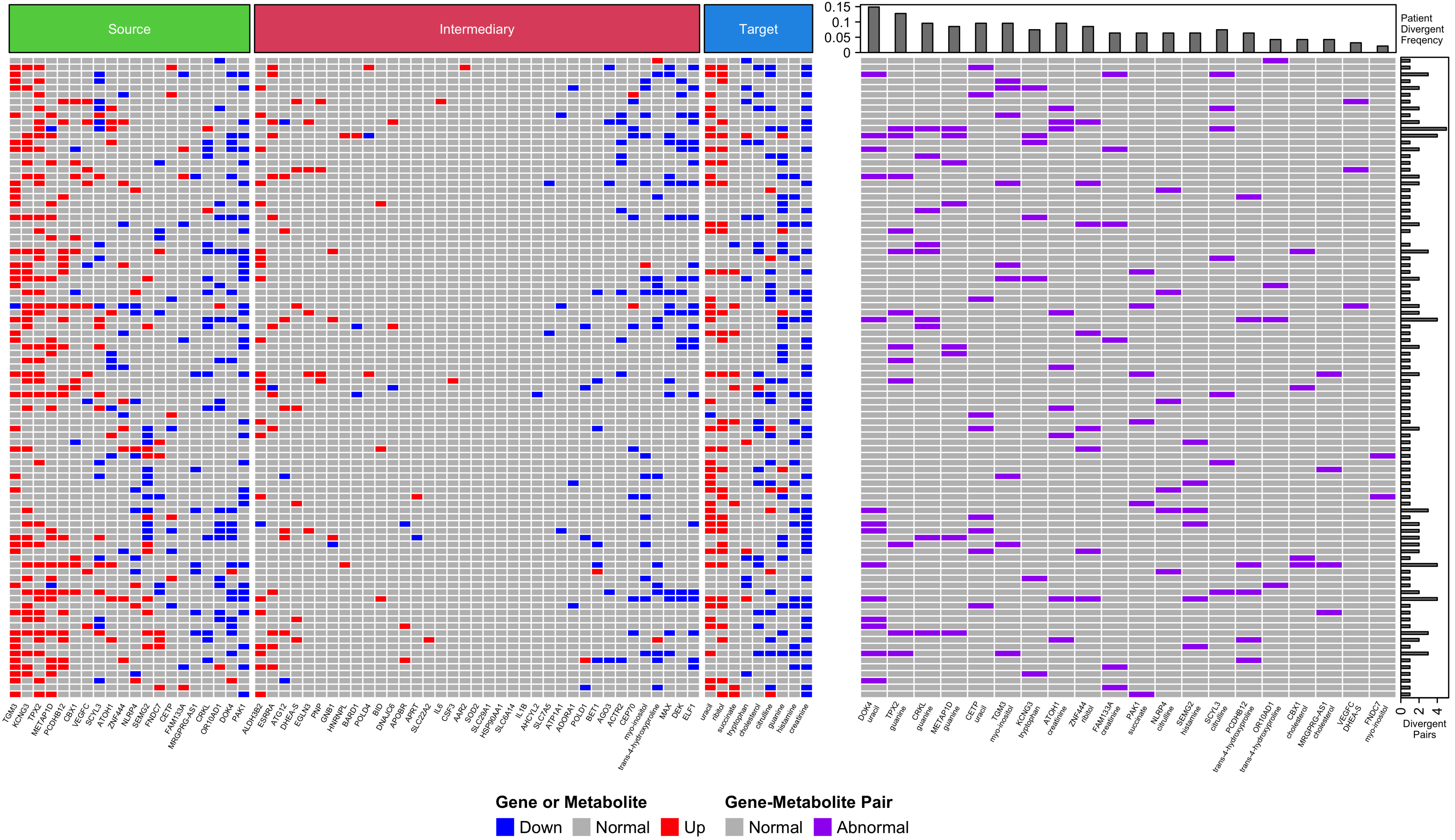

### Figure S2

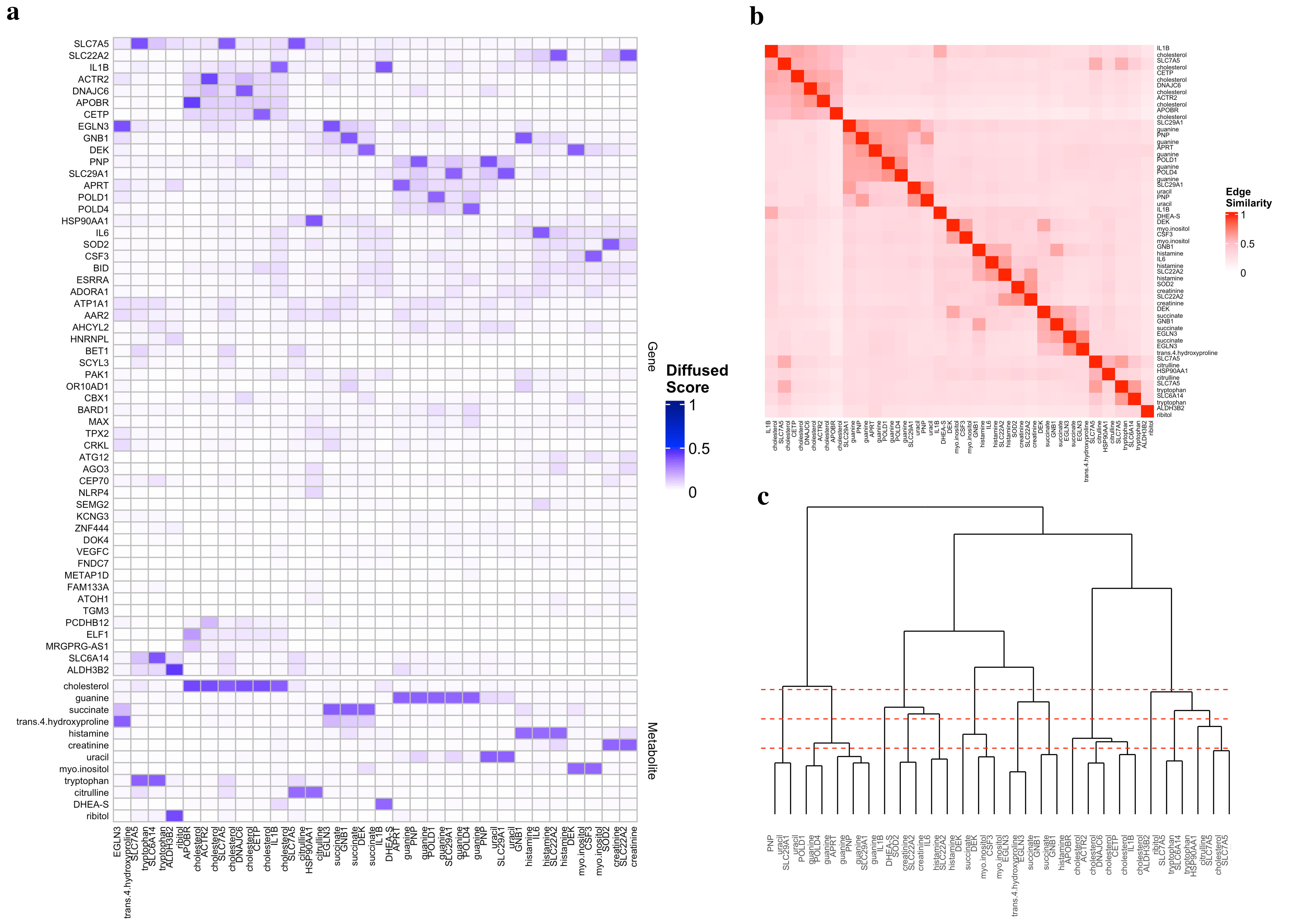
